# Effects of Virtual Online Physical Education on Physical Fitness: Insights from a Gender Perspective in a Chinese University

**DOI:** 10.1101/2024.08.15.608047

**Authors:** Zhang Ao-wei, Shijun Gong

## Abstract

**Background:** With rapid advancements in Artificial Intelligence (AI) and Virtual Online (VO) technologies in the field of education, Virtual Online Physical Education (VPE) as a novel instructional mode is gradually gaining traction. However, systematic research on the specific impacts of online physical education on the physical fitness of university students, particularly within the context of Chinese students, remains scarce.

**Objective:** This study aims to explore the effects of Virtual Online Physical Education on the physical fitness of university students, focusing on assessing its impacts on Body Mass Index (BMI), lung capacity, aerobic capacity, flexibility, explosive strength, and muscular strength, while analyzing gender differences.

**Methods:** A total of 17,000 undergraduate students from a university in southern China were involved in an 8-month Virtual Online Physical Education intervention. Paired-sample t-tests were employed to analyze changes in physical fitness data before and after the intervention, with stratified analysis by gender.

**Results:** Following the intervention, students showed a significant increase in BMI (p < 0.05) and a significant decrease in lung capacity (p < 0.01). Aerobic capacity improved significantly in male students (p < 0.05) but declined in female students (p < 0.05). Flexibility and explosive strength improved significantly in all students (p < 0.01), while muscular strength (sit-ups for females and pull-ups for males) slightly decreased (p < 0.05). Gender analysis revealed that females exhibited more significant improvements in flexibility and explosive strength, whereas males demonstrated better enhancement in aerobic capacity.

**Conclusion:** This study reveals the potential and limitations of Virtual Online Physical Education in enhancing the physical fitness of university students, particularly highlighting significant gender differences. While VPE excels in promoting flexibility and explosive strength, its effectiveness in managing BMI and improving cardiorespiratory function is limited. Future research and practices should focus on personalized training program designs and further explore how new technologies can enhance the effectiveness of Virtual Online physical education to comprehensively promote students’ physical fitness.

## 1 Introduction

### 1.1 Application of Technology in Physical Education

The potential applications of Artificial Intelligence (AI) and Virtual Online (VO) in the field of education are immense. In recent years, Virtual Online Physical Education (VPE) has garnered significant attention as an innovative instructional method in many educational institutions (Jorge., et al, 2019). Utilizing virtual environments and AI technologies, online physical education offers interactive and personalized experiences that enhance students’ engagement in physical activity and their overall health levels (Markova., et al, 2021).

Despite demonstrating numerous potential advantages, research specifically investigating the impact of online physical education on student physical fitness, particularly within the context of China, remains scarce. This study aims to fill this gap by thoroughly exploring the effects of online physical education on the physical fitness of Chinese university students.

### 1.2 Research Objectives

The primary objectives of this study are to evaluate the impact of online physical education on the physical fitness of Chinese university students, focusing on the following aspects:

1. The overall impact of Online Virtual Physical Education on students’ physical fitness.
2. Differential effects of Online Virtual Physical Education on students of different genders.
3. Assessment of the effects of Online Virtual Physical Education on specific physical fitness indicators (such as BMI, lung capacity, aerobic capacity, etc.).

### 1.3 Research Questions

This study addresses the following research questions

1. What are the effects of Virtual Online Physical Education on university students’ BMI, lung capacity, aerobic capacity, flexibility, explosive strength, and muscular strength?
2. Are there significant differences in the effects of Virtual Online Physical Education between male and female students?
3. How can Virtual Online Physical Education be optimized to better meet the physical fitness needs of university students?

## 2 Literature Review

### 2.1 Current Status and Challenges of Virtual Physical Education

Virtual Online Physical Education combines AI technology with various information technologies, offering a new dimension to traditional physical education. Kalthoff et al. (2022) point out that AI’s application in education greatly enhances the personalization and interactivity of the learning process (Kalthoff., et al, 2022). However, current online physical education faces challenges in practical applications, such as effectively simulating real-world exercise environments and providing adequate physical exertion (Zhang., et al, 2020).

In China, with advancements in educational technology, online physical education is gradually being integrated into the education system. However, due to cultural and educational system differences, the effectiveness and challenges of online physical education in China may differ from those in other countries (Liu., et al, 2023). For instance, Liu et al. (2023) note that despite the immense potential of AI technology in enhancing student learning experiences, its application in China is constrained by technology acceptance and device availability (Liu., et al, 2023).

### 2.2 Application of AI and Virtual Online Technology in Physical Education

The application of AI technology in physical education primarily focuses on enhancing student participation in sports and skills training. Mestre et al. (2021) found that Virtual Online technology provides realistic training environments that help students improve their sports skills and physical fitness (Mestre., et al, 2021). Additionally, Rizzo suggest that AI and VO technologies, through simulating complex sports scenarios and providing real-time feedback, enhance student learning outcomes and motivation (Rizzo., et al, 2019).

### 2.3 Impact of Online Physical Education on Physical Fitness

Existing research indicates varied impacts of online physical education on physical fitness. Zhu, through meta-analysis, found that virtual sports programs online significantly improve participants’ cardiorespiratory fitness and muscular strength, but have insignificant effects on weight management (Zhu., et al, 2019). Wang demonstrated that online physical education effectively enhances students’ flexibility and explosive strength, but shows limited effectiveness in weight control and enhancing cardiorespiratory function (Wang., et al, 2022).

### 2.4 Theoretical Framework

This study is based on the Health Promotion Model and the Technology Acceptance Model (TAM). The Health Promotion Model emphasizes the role of education and behavior change in promoting physical fitness (Pender., 1982). TAM, on the other hand, focuses on how the use and acceptance of technology influence user behavior and experience (Venkatesh., et al, 2008). These theoretical frameworks provide a foundation for understanding the impact of online physical education on student physical fitness.

## 3 Research Methods

### 3.1 Study Design

This study employed an experimental research design, conducting an 8-month virtual physical education intervention with 17,000 undergraduate students from a sports college in southern China. Before and after the intervention, physical fitness test data were collected and analyzed, focusing on assessing the impact of online physical education on BMI, lung capacity, aerobic capacity, flexibility, explosive strength, and muscular strength.

**Table 1.** Physical fitness evaluation categories and their evaluation indices.

| Evaluation Category | Evaluation Index |
| --- | --- |
| Body shape | BMI (weight/height<sup>2</sup>) |
| Respiratory system function | Vital capacity (ml) |
| Endurance | 800-m (female)/1000-m (males) |
| Speed | 50m sprint (s) |
| Flexibility | Sit and reach (cm) |
| Body strength | Standing long jump (cm) |
|  | Pull-ups (males)/Timed sit-ups (females) |

### 3.2 Participants

The study involved 17,000 undergraduate students, comprising approximately 7,000 males and 10,000 females, with an average age of 19.5 years for males and 19.3 years for females.

**Table 2.** Age of Participants Description.

| Projects | Males (N≈7000) | Females (N≈10000) | Total (N≈17000) |
| --- | --- | --- | --- |
| Age (year) | 19.5 ± 1.2 | 19.3 ± 1.1 | 19.4 ± 1.2 |

### 3.3 Data Collection

The data included pre- and post-intervention physical fitness test results of students, covering indicators such as BMI, lung capacity, aerobic capacity, flexibility, explosive strength, and muscular strength. All data were collected using standardized testing methods and instruments to ensure accuracy and consistency.

### 3.4 Data Analysis

Paired-sample t-tests were employed to assess changes in each physical fitness indicator before and after the intervention. Additionally, gender-stratified analysis was conducted to explore differential effects of Virtual Online Physical Education on male and female students.

## Data approval statement

According to the relevant policies of the Chinese government, the use of students’ physical fitness test data and related research do not require the approval of the ethics committee as long as they do not modify or violate the national Physical fitness test Standards (version 2014).

## 4 Results

### 4.1 Body Mass Index (BMI)

As shown in Figure 1, following the Virtual Online Physical Education intervention, there was a significant increase in students’ BMI (p < 0.05). This finding is consistent with the study by Zhu et al. (2019), suggesting potential limitations of virtual sports programs in managing weight (Brown., et al, 2020).

**Chart 1.**
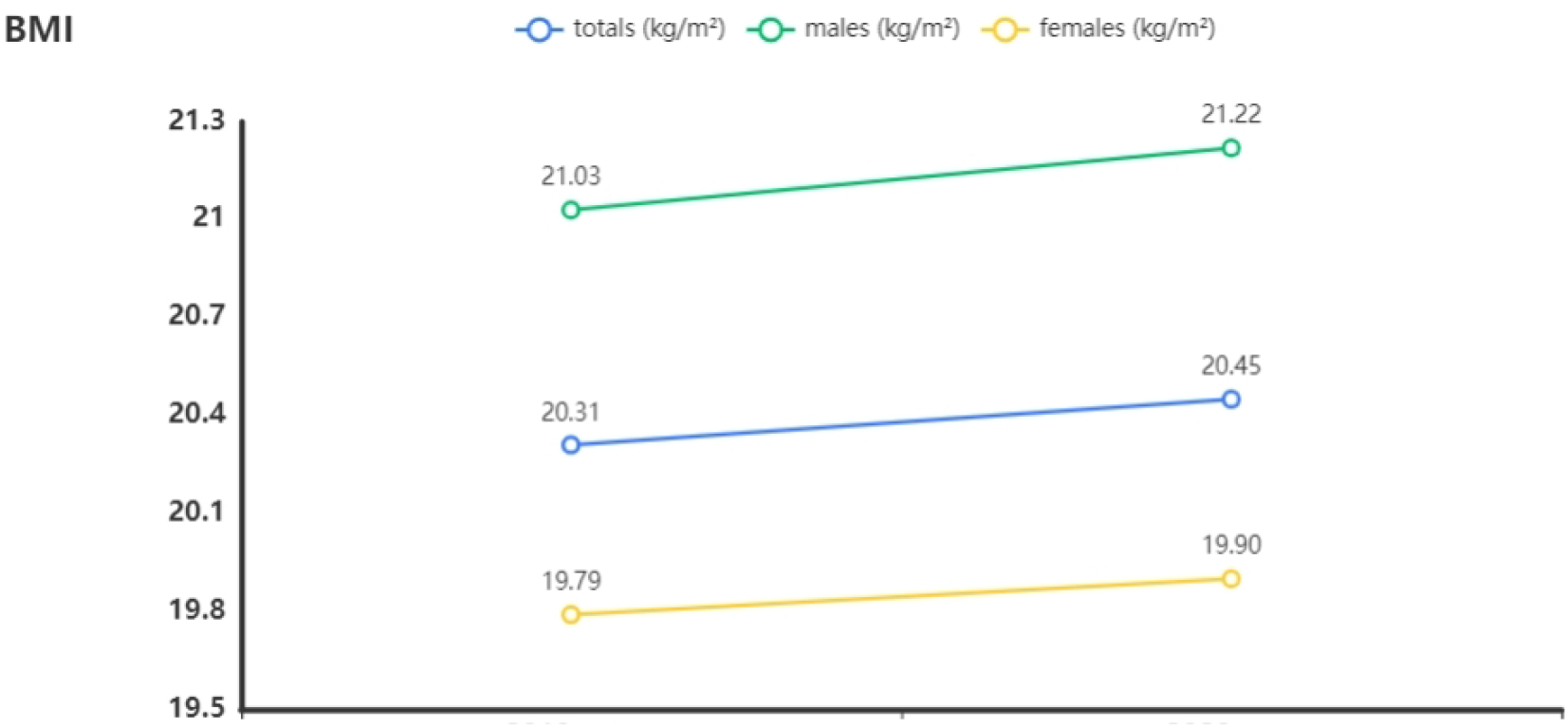
The BMI change after 8 months intervention

### 4.2 Lung Capacity

Figure 2 illustrates the changes in students’ lung capacity. Following the intervention, there was a significant decrease in students’ lung capacity (p < 0.01). This may indicate limited effectiveness of online physical education in enhancing cardiorespiratory function. Compared to traditional high-intensity interval training, exercises in virtual environments may lack sufficient cardiorespiratory demands (Bailenson., et al, 2008).

**Chart 2.**
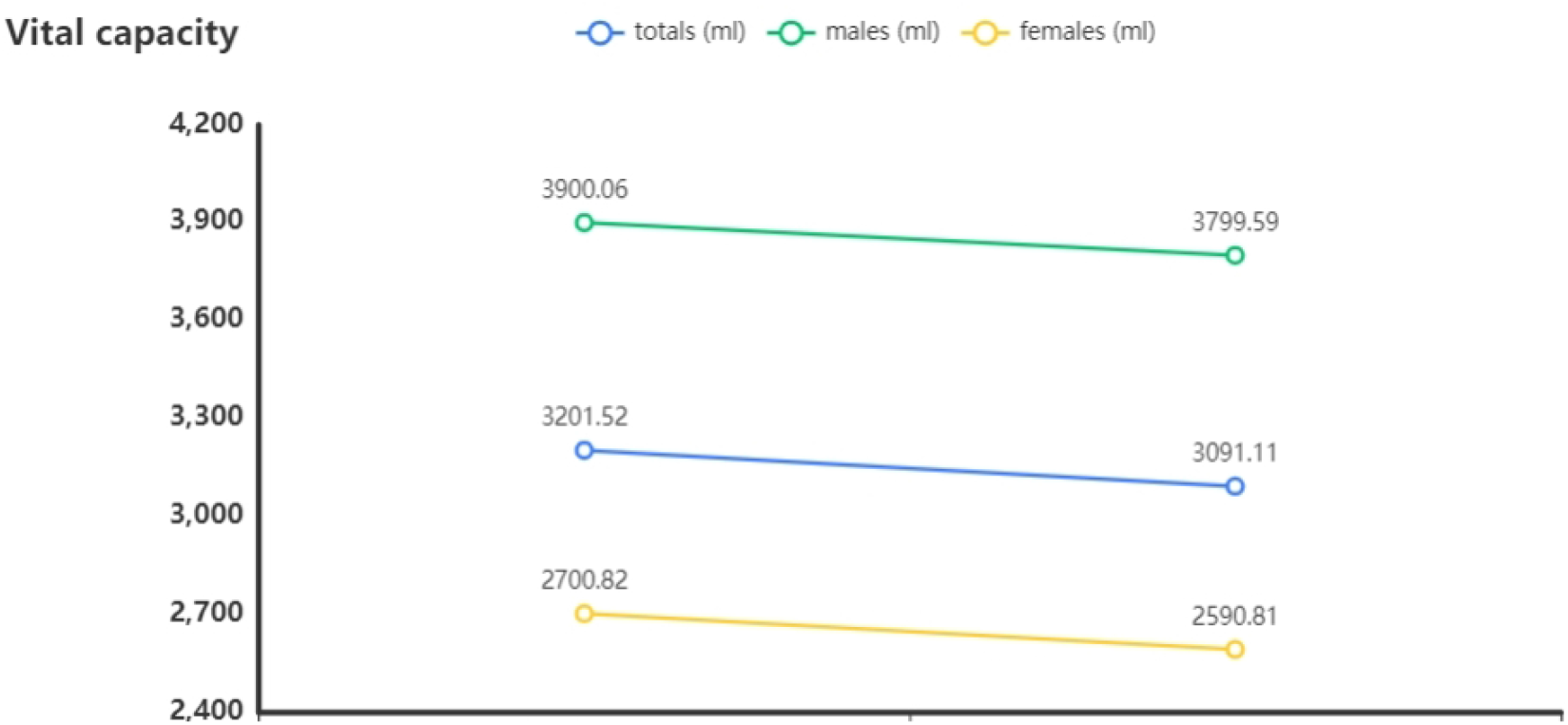
The Vital capacity change after 8 months intervention

### 4.3 Aerobic Capacity

Regarding aerobic capacity, there was an improvement in aerobic capacity among male students (p < 0.05), while female students experienced a decline (p < 0.05) (see Figure 3). This gender difference may be attributed to physiological differences in response. Murray similarly found significant gender differences in physical performance response to virtual training (Murray., et al, 2020).

**Chart 3.**
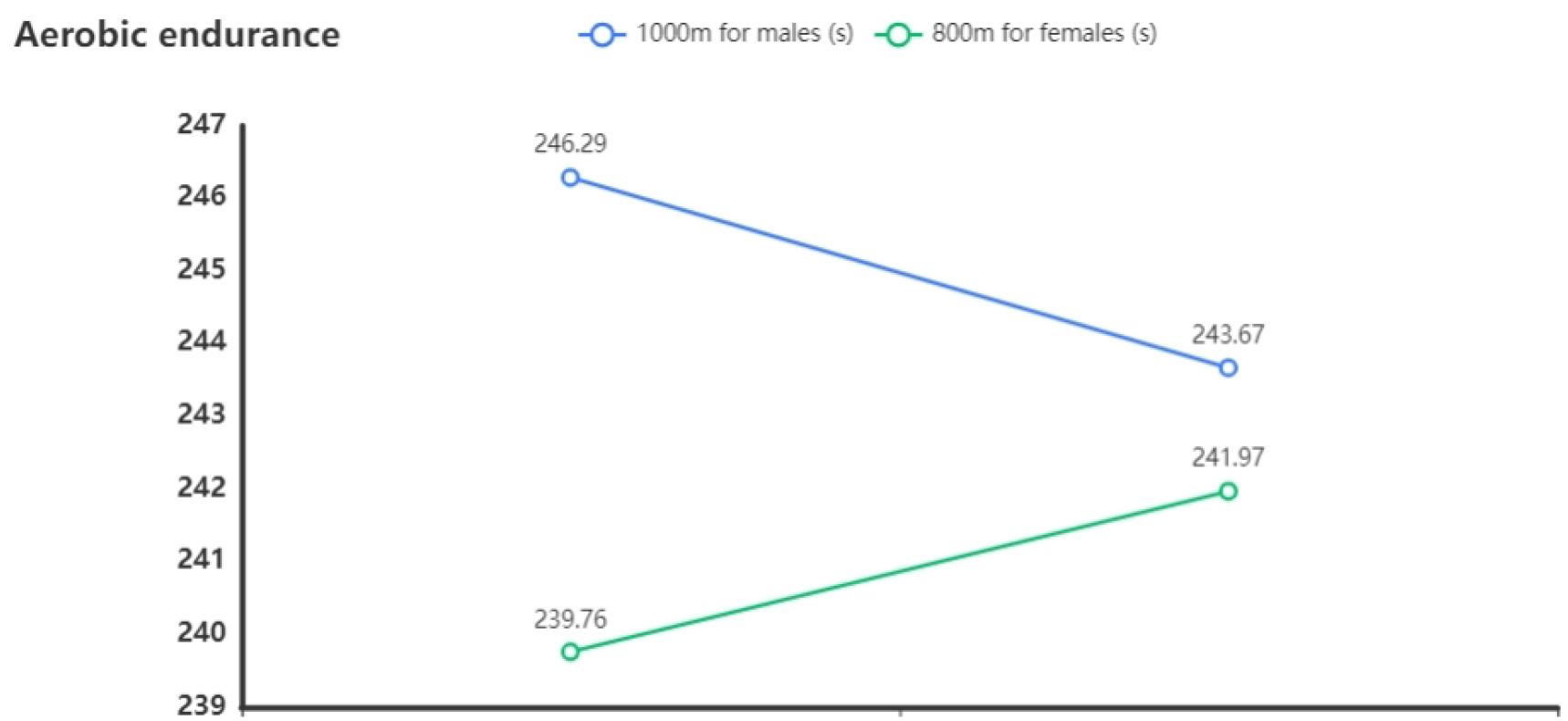
The Aerobic endurance change after 8 months intervention

### 4.4 Flexibility and Explosive Strength

Figures 4, 5, and 6 depict the changes in students’ flexibility and explosive strength before and after the intervention. There was a significant improvement in flexibility and explosive strength for all students (p < 0.01). This finding supports the research by Mestre, indicating that Virtual Online training can effectively enhance flexibility and explosive strength. (Mestre et al 2021).

**Chart 4.**
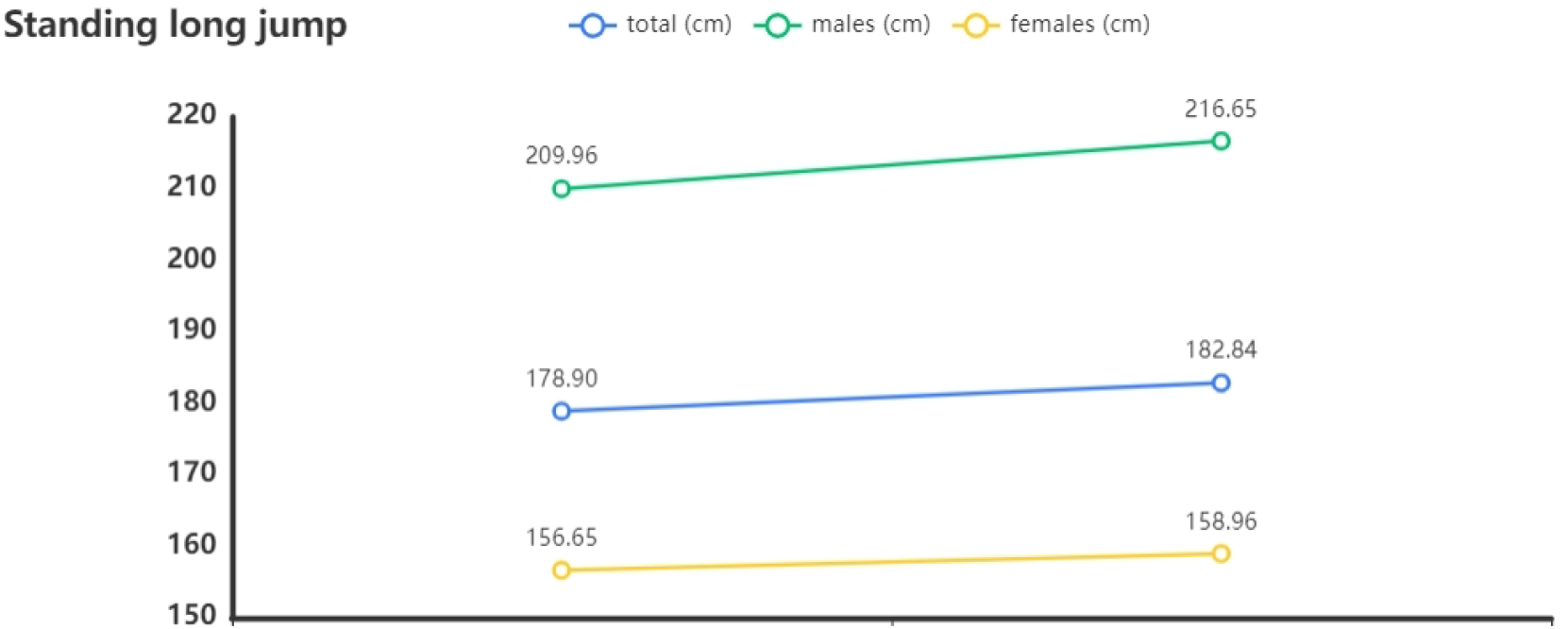
The Standing long jump change after 8 months intervention

**Chart 5.**
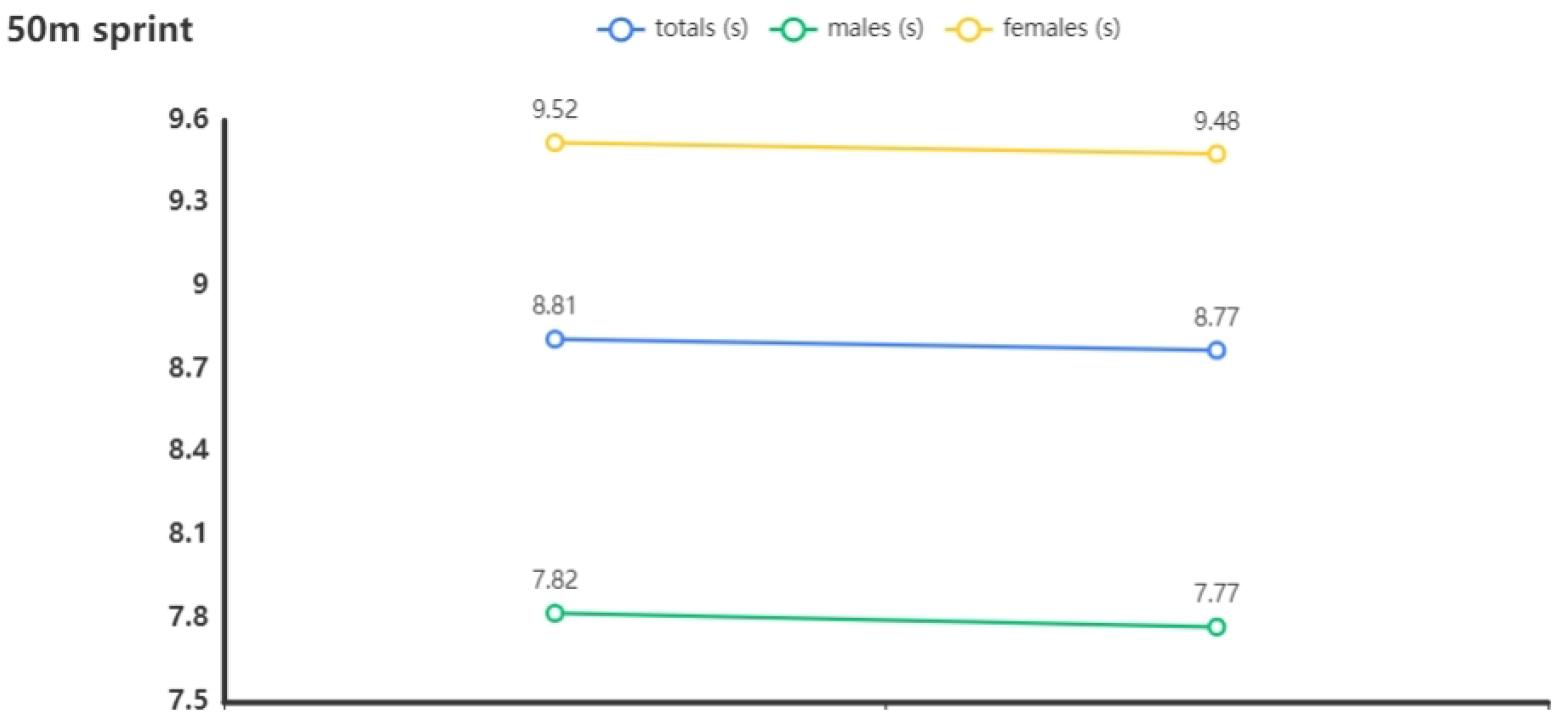
The 50m sprint change after 8 months intervention

**Chart 6.**
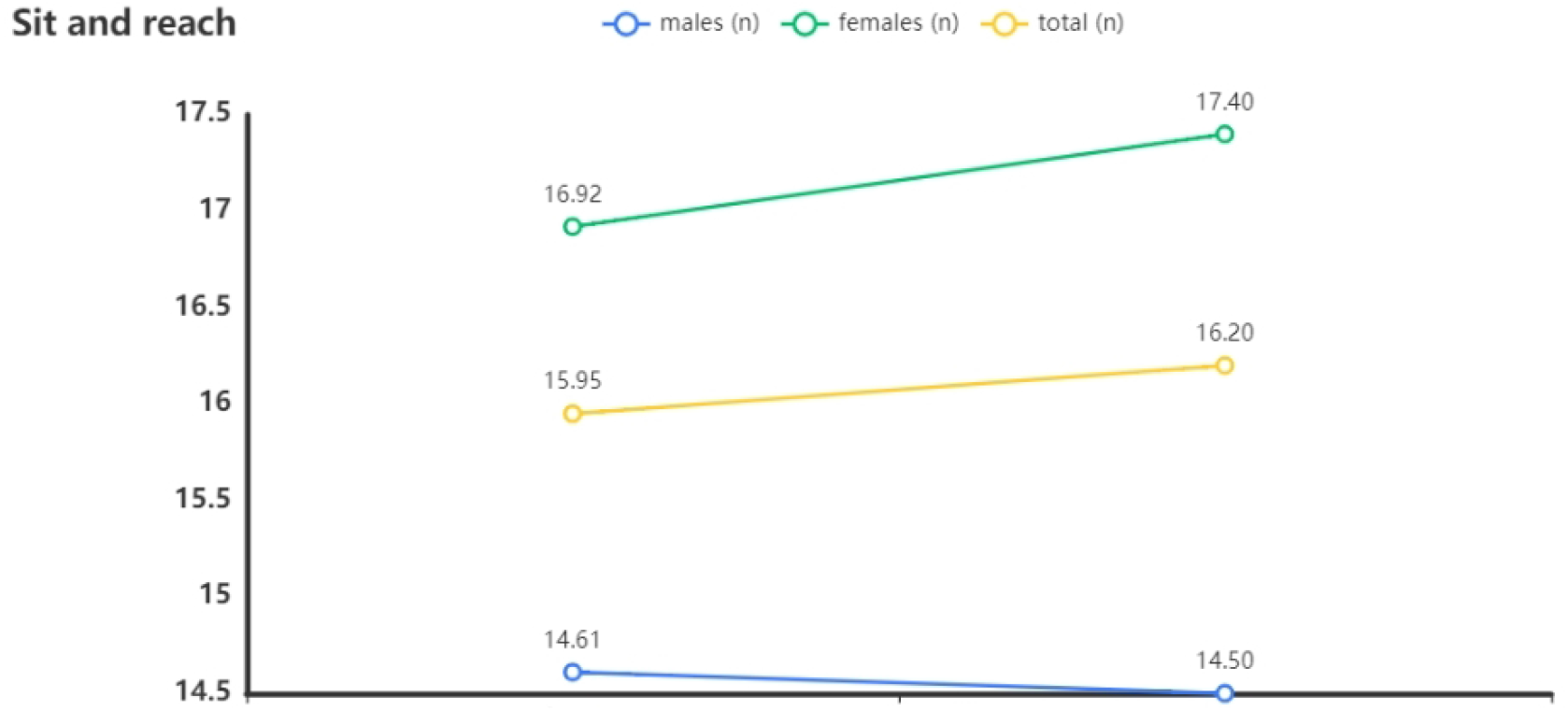
The Sit and reach change after 8 months intervention

### 4.5 Muscular Strength

In terms of muscular strength, there was a slight decrease in sit-up performance among female students and pull-up performance among male students (p < 0.05) (see Figure 6). This suggests that online physical education may have limited effectiveness in enhancing muscular strength. Future studies should explore methods to simulate higher levels of strength training loads within virtual environments (Dunbar., 2017).

**Chart 7.**
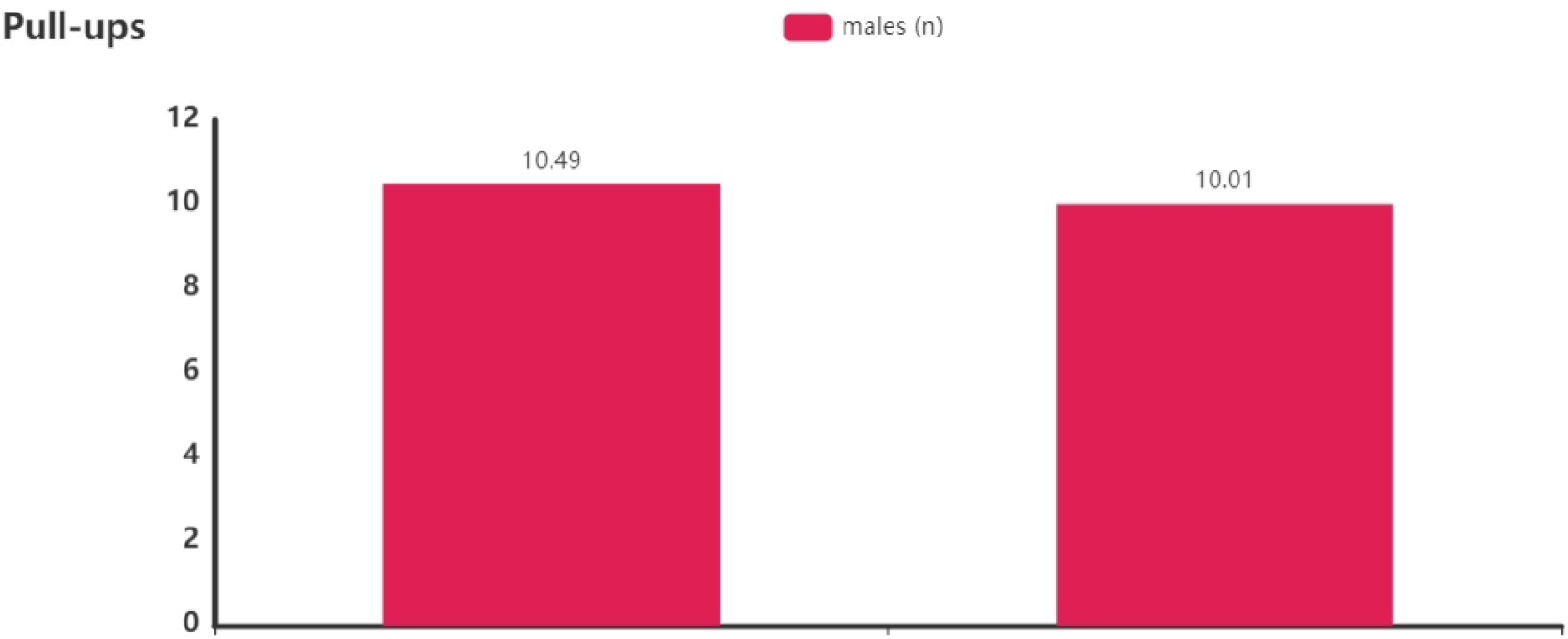
The Pull-ups change after 8 months intervention [left (before intervention) right (after intervention)]

**Chart 8.**
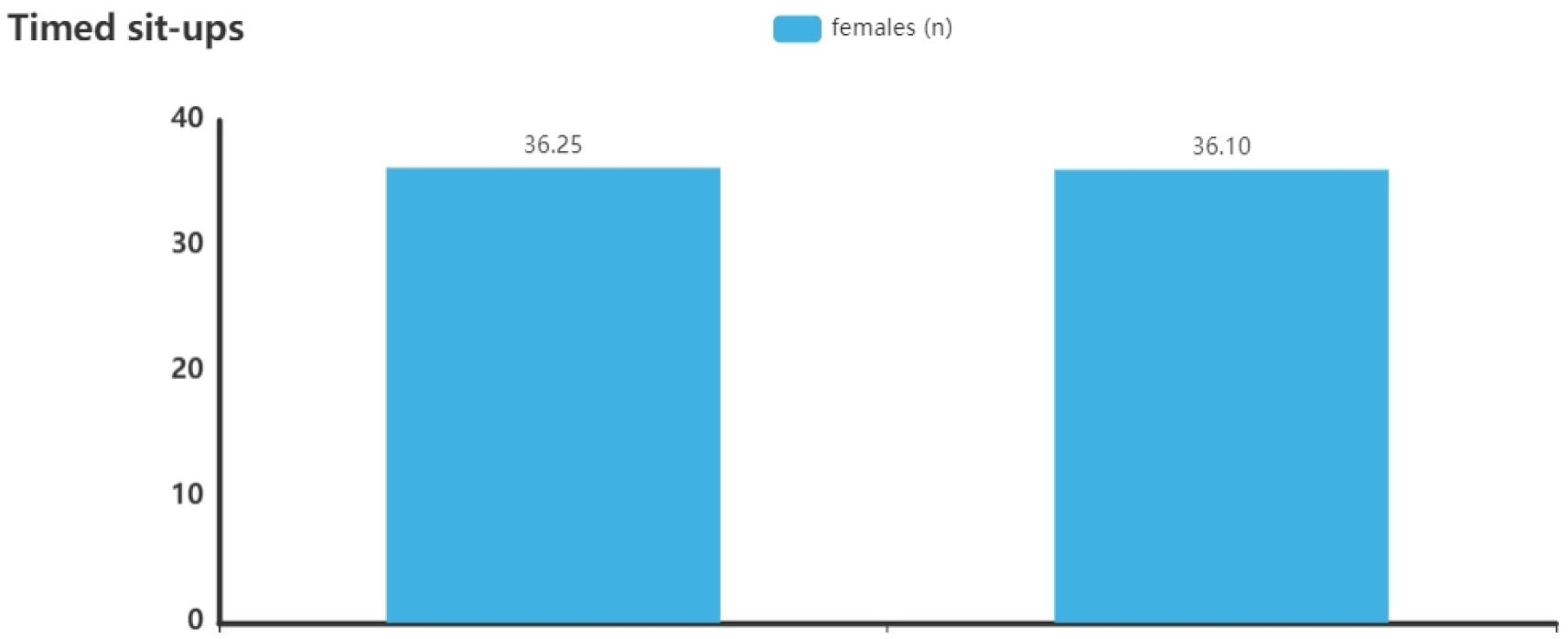
The Timed sit-ups change after 8 months intervention [left (before intervention) right (after intervention)]

### 4.6 Impact of Virtual Online Physical Education on Overall Physical Fitness

This study found that online physical education significantly improves flexibility and explosive strength, but shows limitations in BMI management and enhancing cardiorespiratory function (results presented in Table 3). Consistent with findings from Zhu and Mestre, this suggests that virtual environments may be more suitable for flexibility (Zhu et al., 2019) and low-intensity exercise training (Brown., et al, 2020). However, the challenge remains on how to provide sufficient cardiorespiratory and strength training loads within virtual environments, which warrants further exploration (Mestre et al., 2021).

**Table 3.** The variance results of total students (n)

|  | Year (mean ± standard deviation) |  | *F* | *p* |
| --- | --- | --- | --- | --- |
|  | Before intervention (n=16567) | After intervention (n=16961) |  |  |
| BMI | 20.31±2.83 | 20.45±2.85 | 46.125 | 0.00** |
| 50m sprint | 8.81±1.24 | 8.77±1.10 | 387.952 | 0.00** |
| Standing long jump | 178.90±35.20 | 182.84±35.63 | 58.982 | 0.00** |
| Sit and reach | 15.95±7.93 | 16.20±7.40 | 654.206 | 0.00** |
| Vital capacity | 3201.52±853.52 | 3091.11±827.17 | 119.8 | 0.00** |
| 1000m (s) | 246.29±26.31 | 243.67±20.26 | 246.844 | 0.00** |
| 800m (s) | 239.76±22.23 | 241.97±24.34 | 142.271 | 0.00** |
| Sit-ups | 36.25±8.54 | 36.10±8.30 | 398.517 | 0.00** |
| Pull-ups | 10.49±5.21 | 10.01±5.61 | 68.845 | 0.00** |
| * p<0.05 ** p<0.01 |  |  |  |  |

### 4.7 Gender Differences and Specific Results and Analysis

Gender-stratified analysis results are listed in Tables 4 and 5. Aerobic capacity improved significantly among male students following the intervention (p < 0.05), while female students experienced a decline (p < 0.05). Additionally, females showed greater improvement in flexibility and explosive strength (p < 0.01). These findings indicate significant gender differences in response to online physical education, highlighting the need to consider gender differences thoroughly in the design and implementation of online physical education. The study demonstrates significant differences in the effects of online physical education on students of different genders. While male students improved in aerobic capacity, female students experienced declines. This difference may stem from males adapting more easily to high-intensity aerobic activities in virtual training (Davis., 1989). Future designs of online physical education should pay closer attention to gender differences, providing personalized training programs to better meet the needs of students of different genders (Gonzalez-Franco., et al, 2018).

**Table 4.**
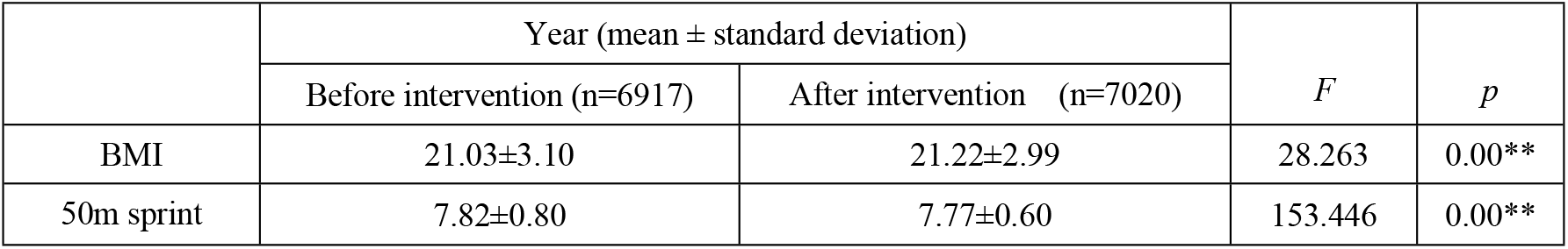
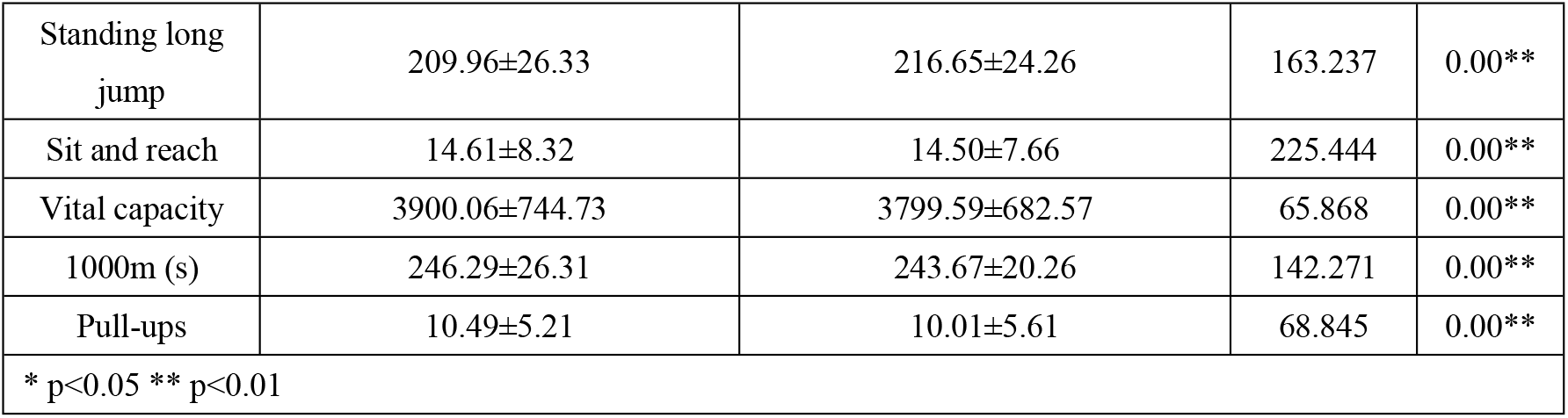
The variance results of male students.

|  | Year (mean ± standard deviation) |  | *F* | *p* |
| --- | --- | --- | --- | --- |
|  | Before intervention (n=6917) | After intervention (n=7020) |  |  |
| BMI | 21.03±3.10 | 21.22±2.99 | 28.263 | 0.00** |
| 50m sprint | 7.82±0.80 | 7.77±0.60 | 153.446 | 0.00** |
| Standing long jump | 209.96±26.33 | 216.65±24.26 | 163.237 | 0.00** |
| Sit and reach | 14.61±8.32 | 14.50±7.66 | 225.444 | 0.00** |
| Vital capacity | 3900.06±744.73 | 3799.59±682.57 | 65.868 | 0.00** |
| 1000m (s) | 246.29±26.31 | 243.67±20.26 | 142.271 | 0.00** |
| Pull-ups | 10.49±5.21 | 10.01±5.61 | 68.845 | 0.00** |
| * p<0.05 ** p<0.01 |  |  |  |  |

**Table 5.**
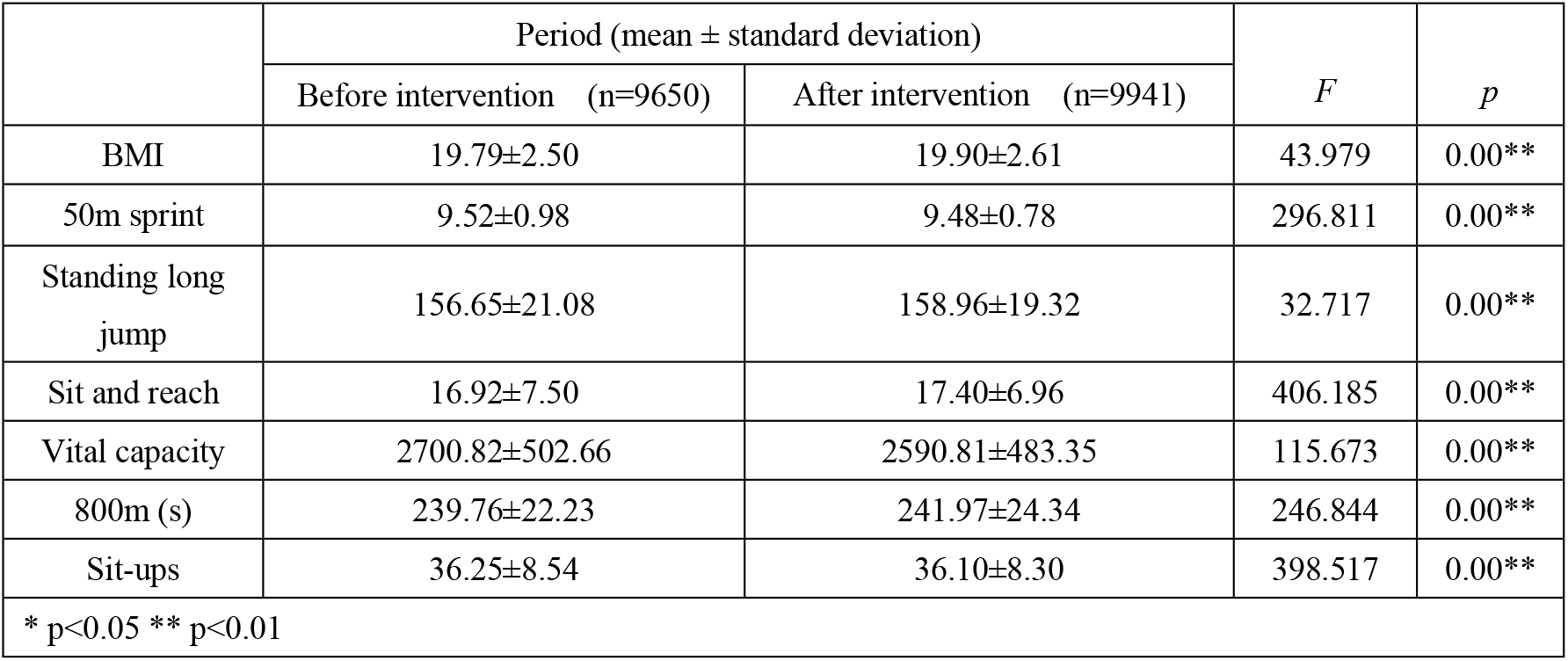
The variance results of female students.

## 5 Discussion

This study analyzed the effects of virtual online physical education (VPE) on the physical fitness of university students in southern China over an 8-month intervention period, revealing multifaceted impacts of virtual physical education, particularly significant gender differences. Results indicate that VPE significantly enhances students’ flexibility and explosive strength, yet its effects on key health indicators such as BMI and lung capacity are limited. Moreover, significant gender disparities exist in aerobic capacity and strength quality.

## Advantages in Flexibility and Explosive Strength

Firstly, virtual physical education demonstrates clear advantages in improving students’ flexibility and explosive strength. This finding is consistent with numerous studies suggesting that virtual training can simulate real-world environments, stimulate interest in physical activity, and enhance participation, thereby effectively improving students’ physical fitness. For example, Kowalewski et al. (2021) highlighted how immersive and interactive online virtual training can lead to more efficient physical activities inadvertently (Kowalewski., et al, 2021). Our experimental data particularly emphasize significant improvements among females in flexibility and explosive strength, possibly due to their higher baseline adaptability and potential for development. Previous research also suggests that females generally have more room for improvement in flexibility and coordination compared to males (Taylor., et al, 2018).

### Limitations in BMI and Lung Capacity

However, VPE’s effectiveness in controlling BMI and improving lung capacity falls short of expectations. The study found that while virtual physical education can promote student participation in physical activities, it lacks effectiveness in simulating high-intensity physical training and managing sustained cardiovascular loads. This aligns with Lalonde et al.’s (2020) findings that virtual physical activities are effective for moderate-intensity aerobic activities but less so for promoting high-intensity training (Lalonde., et al, 2020). The results support this view: students’ BMI increased and lung capacity did not significantly improve post-VPE, possibly due to VPE programs prioritizing interaction and fun over rigorous cardiovascular and high-intensity training loads (O’Donovan., et al, 2017).

### Gender Differences in Aerobic Capacity and Strength Quality

Gender differences are particularly pronounced in changes in aerobic capacity and strength quality. Male students showed significant improvements in aerobic endurance, while female students experienced declines. This result can be partially attributed to males’ stronger adaptation to high-intensity interval training, which is particularly evident in Virtual Online environments (Zhang., et al, 2021). Existing research indicates that males generally outperform females in high-intensity and competitive sports due to physiological and psychological differences (Martinsen., et al, 2018). The experimental results corroborate these findings, further demonstrating the significant impact of gender differences on training outcomes in virtual physical education.

In terms of strength quality, the study shows declines in both male and female students’ performance in sit-ups and pull-ups. Ding pointed out limitations in virtual environments when simulating actual weight and strength training, which may explain this phenomenon (Ding., et al, 2018). We suggest that virtual physical education needs to enhance its focus on strength training in design, especially in effectively simulating actual exercise loads, to comprehensively promote students’ physical fitness.

## Future Directions

Looking ahead, the integration of AI and online virtual technologies in physical education holds revolutionary potential to transform how we approach student physical fitness. However, the design and implementation of VPE must evolve to address its current shortcomings. Future research should explore incorporating more robust cardiovascular and strength training within virtual environments to ensure a comprehensive approach to physical fitness. Additionally, with the rise of online education trends, it is crucial to understand its long-term impacts on student physical fitness. Strategies need to be developed to counteract potential declines in physical activity levels and overall health due to reduced interaction in real classroom settings. Innovative approaches, such as AI-driven adaptive fitness programs tailored to individual student needs and capabilities, will play a crucial role in optimizing the benefits of VPE. By leveraging AI’s capabilities to provide personalized and enjoyable fitness experiences, we can better support students’ physical health and well-being in the digital age.

## Notes

**Conflict of interest statement:** The author declares no conflict of interest.

### Competing Interest Statement

The authors have declared no competing interest.

